# Multi-laboratory assessment of reproducibility, qualitative and quantitative performance of SWATH-mass spectrometry

**DOI:** 10.1101/074567

**Authors:** Ben C. Collins, Christie L. Hunter, Yansheng Liu, Birgit Schilling, George Rosenberger, Samuel L. Bader, Daniel W. Chan, Bradford W. Gibson, Anne-Claude Gingras, Jason M. Held, Mio Hirayama-Kurogi, Guixue Hou, Christoph Krisp, Brett Larsen, Liang Lin, Siqi Liu, Mark P. Molloy, Robert L. Moritz, Sumio Ohtsuki, Ralph Schlapbach, Nathalie Selevsek, Stefani N. Thomas, Shin-Cheng Tzeng, Hui Zhang, Ruedi Aebersold

## Abstract

Quantitative proteomics employing mass spectrometry has become an indispensable tool in basic and applied life science research. Methods based on data-dependent acquisition have proved extremely valuable for qualitative proteome analysis but historically have struggled to achieve reproducible quantitative data if large sample cohorts are comparatively analyzed. Targeted proteomics, most commonly implemented as selected reaction monitoring, has emerged as a powerful alternative and succeeded in providing a data independent approach for reproducible quantitative proteomics data but is limited in the number of proteins quantified. SWATH-MS is a recently introduced technique consisting of a data-independent acquisition and a targeted data analysis strategy that aims to maintain the favorable quantitative characteristics (accuracy, sensitivity, specificity) achieved in targeted proteomics but on the scale of thousands of proteins. While previous SWATH-MS studies have shown high intra-lab reproducibility, this has not been evaluated on an inter-lab basis. In this multi-laboratory evaluation study using data from 11 sites worldwide, we have demonstrated that using SWATH-MS we can consistently detect and quantify more than 4,000 proteins from HEK293 cells and that the quantitative protein data generated across laboratories is reproducible. Using synthetic peptide dilution series, we have shown that the sensitivity, dynamic range and reproducibility established with SWATH-MS methods are also uniformly achieved across labs. This study demonstrates that SWATH-MS is a reproducible and accurate technique that can be confidently deployed for large-scale protein quantification in life science research.

## Introduction

Reproducibility is an essential foundation of scientific research. Recent reports have concluded that a significant fraction of life science research shows poor reproducibility of results and this poses a major challenge to scientists, science policy makers, funding agencies and the pharma and biotech industry sectors^1–3^. The reasons for irreproducibility of research results are many, including inadequate study design and data analysis, limited data quality, incompletely characterized research reagents, poorly benchmarked techniques, and a range of other confounding factors.

The question of whether specific data acquisition/data analysis methods and platforms are capable of generating reproducible results is best addressed by inter-laboratory studies, where samples of known composition and quality are analyzed across different settings. Such studies have been reported for various ‘omics’ technologies, including RNA-seq and microarray techniques, with varying results ^4,5^. Such projects have served to highlight problems in various large-scale strategies, to stimulate discussion in a given field on how to improve reproducibility, and in the best cases to provide confidence in a given strategy within and beyond an analytical field.

In the field of mass spectrometry (MS) based proteomics, a wide range of specific methods have been reported over the past two decades. These can be broadly grouped into discovery and targeted proteomic techniques. The general aim of discovery proteomics is the identification of the protein components of biological samples. This is most frequently achieved by data-dependent acquisition (DDA) whereby a fragment ion spectrum is generated from each sequentially selected precursor ion and then associated with the best matching peptide sequence by database searching, followed by inference of a set of protein identities. If the number of detected precursor ions significantly exceeds the number of precursor selection cycles^6^, precursor selection becomes a stochastic process and the peptides detected in repeat analyses of the same sample become irreproducible. This has been documented in a number of intra-and inter-laboratory studies^7–9^. In general, these studies confirmed that a high degree of reproducibility is difficult to achieve for complex samples, even if substantial measures are implemented to control experimental variables^10^. Computational methods to enable improved quantification via propagation of peptide identifications across runs via alignment of MS1 precursor signals, first introduced as accurate mass and time tags (AMT)^11,12^, are commonly applied to DDA data^13–16^ and can reduce this issue to some degree in discrete datasets where chromatographic alignment can reasonably be applied.

In contrast to discovery proteomics the general aim of targeted proteomics is the detection and quantification of a predetermined set of peptides in a sample by selected reaction monitoring (SRM) also known as multiple reaction monitoring (MRM)^17^, or a related technique parallel reaction monitoring (PRM)^18–20^ These analyses require the prior generation of a specific measurement assay^21,22^ for each targeted peptide. The coordinates that constitute the assay frequently include information such as the *m/z* of precursor and fragment ion pairs, their relative intensity, and normalized chromatographic retention time. The coordinates are used to direct the instrument to selectively acquire signals corresponding to the targeted peptides over time, and to confirm the presence of each peptide by comparison of the acquired data to the reference information. Because targeted MS eliminates the stochastic component of precursor ion selection in DDA, it has the potential for high reproducibility. This has been demonstrated in intra-laboratory studies where sets of peptides were targeted with a high degree of reproducibility across relatively large sample sets^23–25^ and by inter-laboratory studies focused on exploring the use of SRM and immuno-SRM for biomarker studies^26–31^. Targeted MS is now broadly regarded as a reproducible protein analysis platform^17^. However, the number of proteins measured is restricted (usually to ~100 per injection), thus limiting its utility as a proteomic technique for many applications.

SWATH-MS is a more recently introduced approach to MS-based proteomics^32^. It consists of data-independent acquisition (DIA) in which all precursor ions within a user defined *m/z* window are deterministically fragmented. Analysis of SWATH-MS data most often relies on a targeted data analysis strategy in which target peptides are detected and quantified from the SWATH-MS fragmentation data by extracting and correlating previously generated query parameters for each target. In this scheme each unique peptide of interest at a given precursor charge state is queried for in the data, resulting in the detection and scoring of co-eluting transition group signals and associated underlying mass spectral features, referred to as peak groups. Because the method specifically tests for the presence of each target peptide in the essentially complete fragment ion map of each sample, it eliminates the stochastic sampling element of DDA and helpfully provides a direct statistical measure (e.g. q-value) of whether the peptide is present at a detectable level in the sample. This data analysis strategy, whereby target peptides are directly queried for, has recently been generalized using the term peptide-centric^33–34^ scoring to distinguish from more classical approaches where the MS2 spectrum is the query unit for data analysis (referred to as spectrum-centric scoring). The SWATH-MS implementation of the DIA concept therefore preserves the favorable performance characteristics of SRM, while vastly expanding the measurement capacity to thousands of proteins per injection. Of consideration in SWATH-MS is the complexity of the resultant spectra and specific software tools have been compiled to analyze such highly multiplexed data using various approaches^35–38^. A recent study comparing software tools for the analysis of DIA data using either peptide-centric or spectrum centric approaches has demonstrated that very similar qualitative and quantitative results can be obtained when analyzing a benchmarking dataset^39^. SWATH-MS and related DIA approaches have achieved a high degree of reproducibility in intra-laboratory studies in a variety of research questions such as interaction proteomics^40,41^, plasma proteomics^42^, tissue proteomics^43^, microbial proteomics^44,45^, pre-clinical toxicology^9^, analysis of genetic reference strains^46^ and many others. However, interlaboratory robustness and reproducibility has not been demonstrated.

In this study, we set out to test the reproducibility of peptide and inferred protein detection and quantification by SWATH-MS in an inter-laboratory study. To achieve this goal we distributed benchmarking samples to 11 participating laboratories worldwide for measurement by SWATH-MS according to a predetermined schedule. We analyzed the data from all sites centrally with two separate scopes in mind. Firstly, we analyzed all of the data in an aggregated way to simulate, for example, a large cohort study whereby patient samples would be analyzed in multiple laboratories, aiming to achieve a result set based on all samples. In the second interpretation, we analyzed the data from each site of collection independently and we compared the results across sites post-analysis facilitating a direct performance comparison.

Our analysis demonstrated that the set of proteins detected and quantified across all participating sites, i.e. from a total of 229 proteome measurements, was very consistent. The reproducibility, linear dynamic range, and sensitivity are approaching those reported for SRM, currently the gold standard approach for protein quantification^17^. This data supports the conclusion that DIA combined with peptide-centric scoring embodied by the SWATH-MS approach is suitable for both comprehensive and reproducible proteomics at large-scale and across laboratories.

## Results

### Study design and implementation

To assess the inter-and intra-laboratory reproducibility and performance of SWATH-MS for large-scale quantitative proteomics, we created a benchmarking sample set and distributed aliquots to 11 laboratories worldwide (Fig 1a). The sample consisted of 30 stable isotope labeled standard (SIS) peptides^47^ diluted into a complex background consisting of 1μg of protein digest from HEK293 cells. To achieve both, a physiologically-relevant fold change step, and to cover a large linear dynamic range in a relatively small number of samples that could be analyzed in a 24 hour period, we elected to partition the SIS peptides into 5 groups (A-E), each containing 6 peptides. In each group, the dilution series started from a different level ranging from 1 fmol to 10 pmol (sample S5). The peptides were then diluted serially 3-fold into the HEK293 background 4 times (samples S4-S1). This generated an overall dilution series from 0.012 to 10,000 fmol on column, with a linear dynamic range over 6 orders (although not covered by any single SIS peptide) and maintaining a physiologically-relevant fold change of 3 **(Supplementary Tables 1-2).** We acquired all data in SWATH-MS mode, set to 64 variable width Q1 windows chosen to minimize window size in high density precursor ion ranges **(Supplementary Table 13)**.

**Figure 1:**
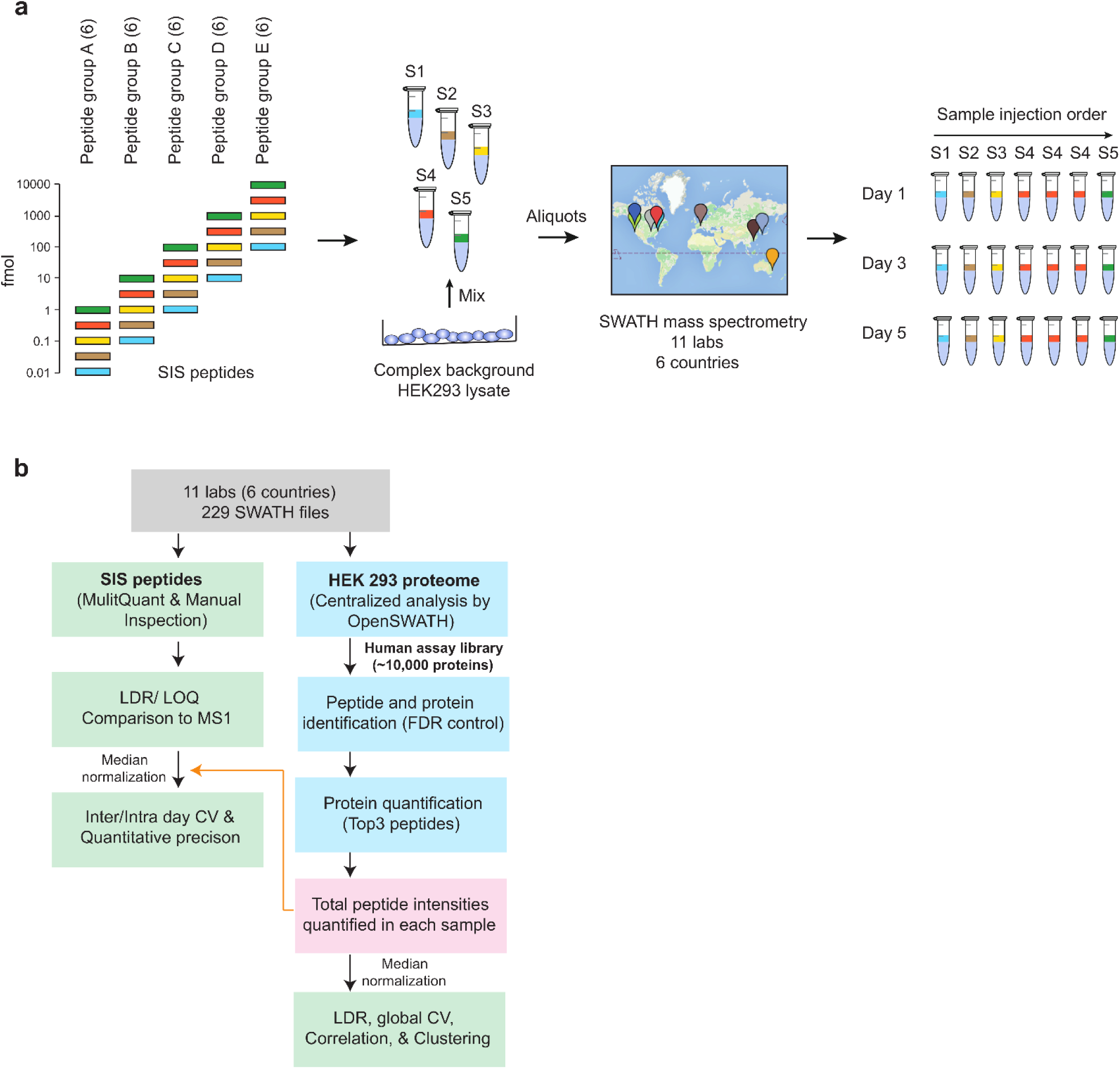
Study design and implementation. (a) A set of 30 SIS peptides partitioned into 5 groups (A-E, 6 peptides in each) were diluted into a HEK293 cell lysate to span a large dynamic range. Starting at a different upper concentration for each group, they were 3-fold diluted into the matrix to cover a concentration range from 12 amol to 10 pmol in lμg of cell lysate. This created a set of 5 samples to be run by SWATH-MS on the TripleTOF 5600/5600+ system at each site. Each sample was run once per day on day 1, 3 and 5, with the exception of sample 4 which was run 3x on each day. (b) After data acquisition, the 229 SWATH-MS files were assembled centrally and processed using two strategies. The SIS peptide concentration curves were assessed using MultiQuant Software, allowing for the determination of linear dynamic range (LDR), and LLOQs for each peptide. In addition, the intra-and inter-day CVs were determined before and after normalization. The HEK293 proteome matrix data was analyzed using the OpenSWATH pipeline and the Combined Human Assay Library consisting of ~10,000 proteins. The false discovery rate was controlled at the peptide query and protein level using PyProphet. Protein abundances were inferred by summing the top 5 most abundant fragment ions from the 3 most abundant peak groups using the aLFQ software. We then used protein abundances to cluster, and compute Pearson correlation coefficients, for all samples from all sites.

To standardize the SWATH-MS acquisition protocol and to make an initial quality assessment we first asked each site to acquire 5 replicate injections of a test sample containing only the HEK293 background. This data was used to improve quality control procedures and to ensure adequate system performance at all sites **(Supplementary Fig. 1; Supplementary note 1).** The finalized study protocol is provided **(Supplementary Protocol 1).** All sites used the same mass spectrometer (SCIEX TripleTOF^®^ 5600 / 5600+ systems), while the nanoLCs consisted of various models from the same vendor (SCIEX). The chromatographic columns had the same dimensions (30cm x 75μm) although 9 sites used cHiPLC^®^ microfluidic systems and 2 sites used self-packed columns with emitters and, as such, therefore also used different chromatographic resins (see **Online Methods; Supplementary Table 11).** After the initial quality control phase, participating labs acquired SWATH-MS data for the main sample set consisting of samples S1-S5 with sample S4 injected in technical triplicates, and repeated this acquisition scheme two further times during 1 week. The purpose of this design was to determine reproducibility and quantification metrics within 1 day, across 1 week, or across different sites of data collection. These measurements resulted in a dataset containing in total 229 SWATH-MS files from the 11 sites worldwide which are freely accessible for further analysis by the community.

### Consistency of protein detection

The qualitative similarity of SWATH-MS data acquired at different sites was investigated by comparing the set of proteins detected from the HEK293 proteome across all 229 SWATH-MS data files. Targeted analysis was performed using the OpenSWATH software^35^ combined with a previously published SWATH-MS spectral library containing peptide query parameters mapping to 10,000+ human proteins^48^ (Fig. 1b). The false discovery rate (FDR) was controlled at 1% at the peptide query and protein levels using the q-value approach^49–53^ in the global context, and at 1% peptide query FDR on a sample-by-sample basis. We did not employ any alignment or transfer of peptide identification confidence between runs. A description of FDR calculation, and issues surrounding this, is provided in **Supplementary Note 2** and in the accompanying manuscript^54^.

The results are shown in Fig. 2. In Fig. 2a we depict the number of proteins detected across all SWATH-MS acquisitions in the aggregated data analysis (equivalent plot at peptide query level in **Supplementary Fig. 2).** The total number of proteins detected at 1% FDR over the entire dataset is 4,984 from 40,304 proteotypic peptide peak groups **(Supplementary Table 3).** The median number of proteins detected per file is 4,710 from a median of 34,480 peak groups. 4,077 proteins were detected in >80% of all samples. Fig. 2b shows the distribution of complete/missing values from this data. Of the 4,984 proteins detected, 3,985 were detected using >1 peptide peak group and, on average, we detected 8.1 proteotypic peptides per protein **(Supplementary Fig. 3).** Information regarding mass spectrometric and chromatographic performance metrics across the sites that might affect the number of proteins detected is provided in **Supplementary Figures 4-8.** The accumulation of new protein identifications over the dataset - indicated by the blue curve in Fig. 2a - saturates, indicating the comprehensiveness of the SWATH-MS methodology and the minimal number of accumulated false positive identifications across 229 measurements. This also indicates that when we analyzed the data in an aggregated manner (i.e. data from all sites combined) the set of proteins detected by all labs is very consistent. Achieving this consistency was dependent on appropriate FDR control in the global context at both peptide query and protein level. To illustrate this, we plotted the numbers of peak groups and proteins detected when FDR was controlled only at peptide query level and not the protein level, and only on a sample-by-sample basis and not in the global context **(Supplementary Fig. 9).** The accumulation of new peak groups steadily increased across the dataset, indicating a likely accumulation of false positives and, highlighting the importance of appropriate global FDR control^54,55^.

**Figure 2:**
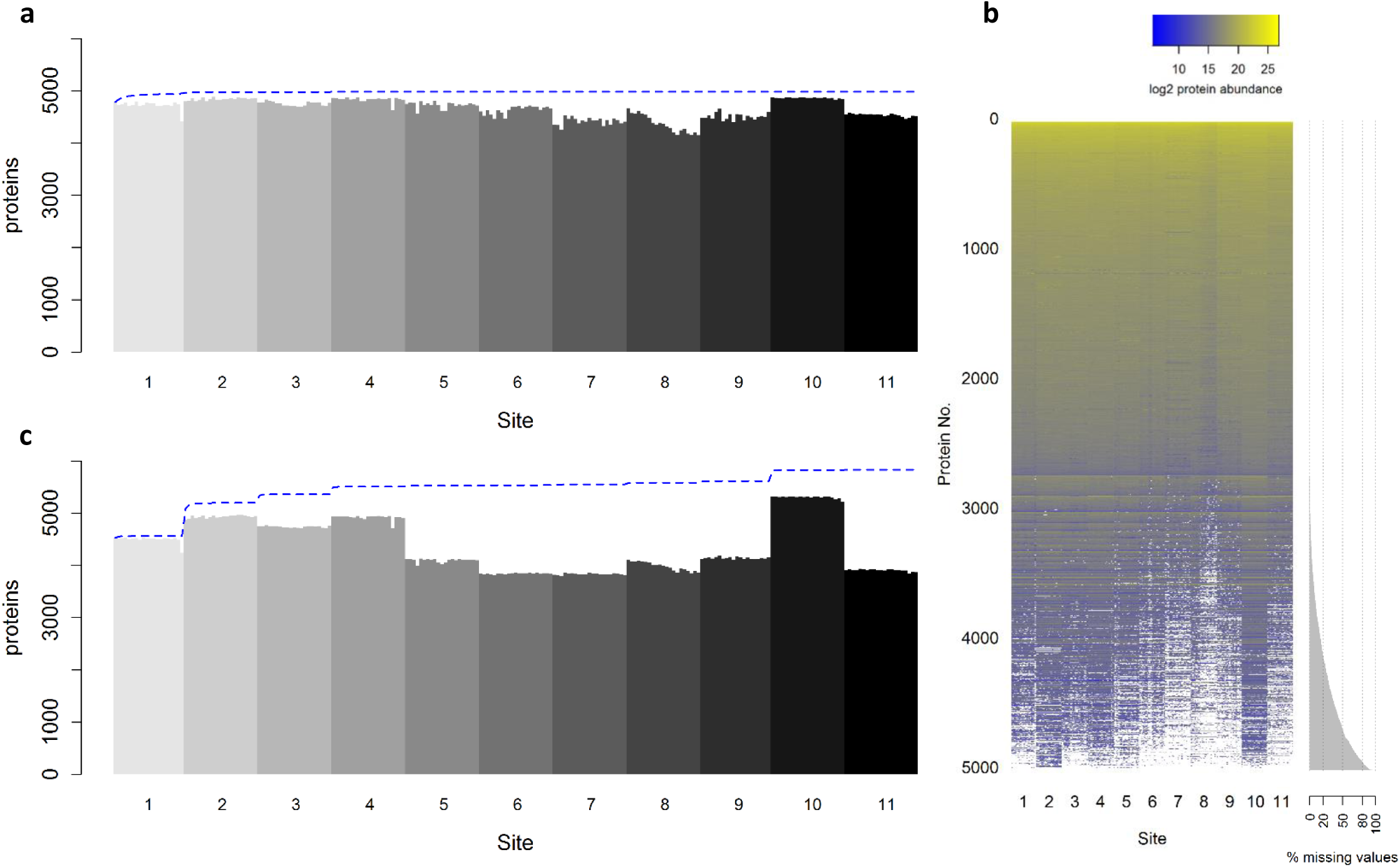
A consistent set of proteins is detected across sites. (a) The number of proteins detected in each of the 229 SWATH-MS analyses is shown ordered by site of data collection and then chronologically by time of acquisition. After filtering the dataset in a global fashion at 1% FDR at the peptide query and protein levels, a protein was considered detected in a given sample when a peak group for that protein was detected at 1% FDR in the context of that sample (see **Supplementary Note 2** for a detailed discussion of FDR). The blue line indicates the cumulate set of proteins detected with each new sample moving from left to right. The maximum of the blue line indicates the set of proteins detected at 1% FDR in the global context. The saturation of the number of proteins detected after a few samples indicates that the set of proteins observed by all sites is highly uniform. (b) A protein abundance matrix on the log2 scale is shown for 229 SWATH-MS analyses from all sites corresponding to the set of proteins shown in panel (a). White indicates a missing protein abundance value where a given protein was not confidently detected in a given sample. The proteins are ordered from top to bottom first by row completeness and then by protein abundance. (c) Equivalent to panel (a) except that the analysis and FDR control is carried independently out on a site-by-site basis instead of aggregated across all sites before analysis and FDR control.

The comparison between the protein detection rates from the aggregated analysis and an individual site-by-site analysis also provides insight into FDR control. Fig. 2c shows the number of proteins detected when the data from each site was first analyzed separately by site of data collection with independent FDR control and then aggregated (equivalent plot at peptide query level in **Supplementary Fig. 10).** In this analysis, the procedure was identical to that of the aggregated analysis, except that the global context for FDR control mentioned above was restricted to the files from an individual site, and that procedure was repeated for each site individually. The information content of the data from each site is not identical, which likely relates to performance differences between chromatographic, nanospray ionization and/or instrument efficiencies across sites at the time of data acquisition. When the data is aggregated before analysis and FDR control, the higher quality data effectively supports the lower quality data, because the strict scoring cutoffs required by the 1% protein FDR threshold only needs to be achieved once per protein in the global context, leading to more homogenous results in terms of proteins detected. That is, in our analysis, a protein is considered detected in a given sample if it is detected at the 1% peptide query FDR threshold as long as the peptide has been detected elsewhere in the experiment with a score passing the 1% protein FDR threshold (see **Supplementary Note 2**).

From these analyses, we can conclude that using SWATH-MS data collected from instruments in different labs, the set of proteins detected is comparable **(Fig 2a and 2b)** when using a data analysis strategy employing peptide-centric scoring. This presents a desirable quality not previously demonstrated at this scale in large-scale proteomics analysis.

### Reproducibility of quantification

Having established a high degree of reproducibility of protein detection within and across sites, we went on to investigate the quantitative characteristics of our inter-lab SWATH-MS dataset. To determine quantitative reproducibility we computed the coefficient of variation (CV) at different levels. Firstly, we extracted ion chromatograms (XIC) for the SIS peptides using the MultiQuant™ software **(Supplementary Table 4).** Next, we computed the CV for each site within 1 day (intra-day) and over the week (inter-day) for the S4 sample, which was acquired every day in triplicate. The median for site intra-day and inter-day CVs (expressed as median ± standard deviation) were 5.5 ± 2.9% and 8.9 ± 11.1%, respectively **(Fig. 3a; Supplementary Table 4 and 5).** For the majority of sites the intra-day and inter-day CVs were below 20% (one lab - Site 8 - experienced some larger LC-MS variance over the course of the week with decreasing signals that was later explained by a contaminated collision cell). As the signal response varies between instruments, attempting to directly compare raw peak area or intensities across sites is not feasible. To determine if we could normalize the instrument response differences by applying a simple normalization, we used the quantitative information from the HEK293 proteome that is expected to be invariant. Specifically, the peptide peak areas from the automated OpenSWATH analysis for each of the 229 files were re-scaled such that the median values from each file were equalized. The resulting protein abundance boxplots in **Supplementary Fig. 11** clearly show the effect of this simple normalization. The normalization coefficients **(Supplementary Fig. 12)** were used to adjust the peptide peak areas for the SIS peptides derived from the MultiQuant analysis and the intra-day and inter-day CV analysis was repeated **(Fig 3a).** We then calculated the inter-site CVs for the SIS peptides using all measurements of S4 sample from all sites. The median of the inter-site CVs using peptide peak areas without normalization was 47.3 ± 13.9%. After normalization, this was reduced to 21.3 ± 10.3%. Normalization also reduced the median within site inter-day CV from 8.9 ± 11.0% to 5.8 ± 5.4% whereas the intra-day CV was less strongly affected (5.5 ± 2.9% to 4.7 ± 2.3% intra-day CV) **(Supplementary Table 4-5).**

**Figure 3:**
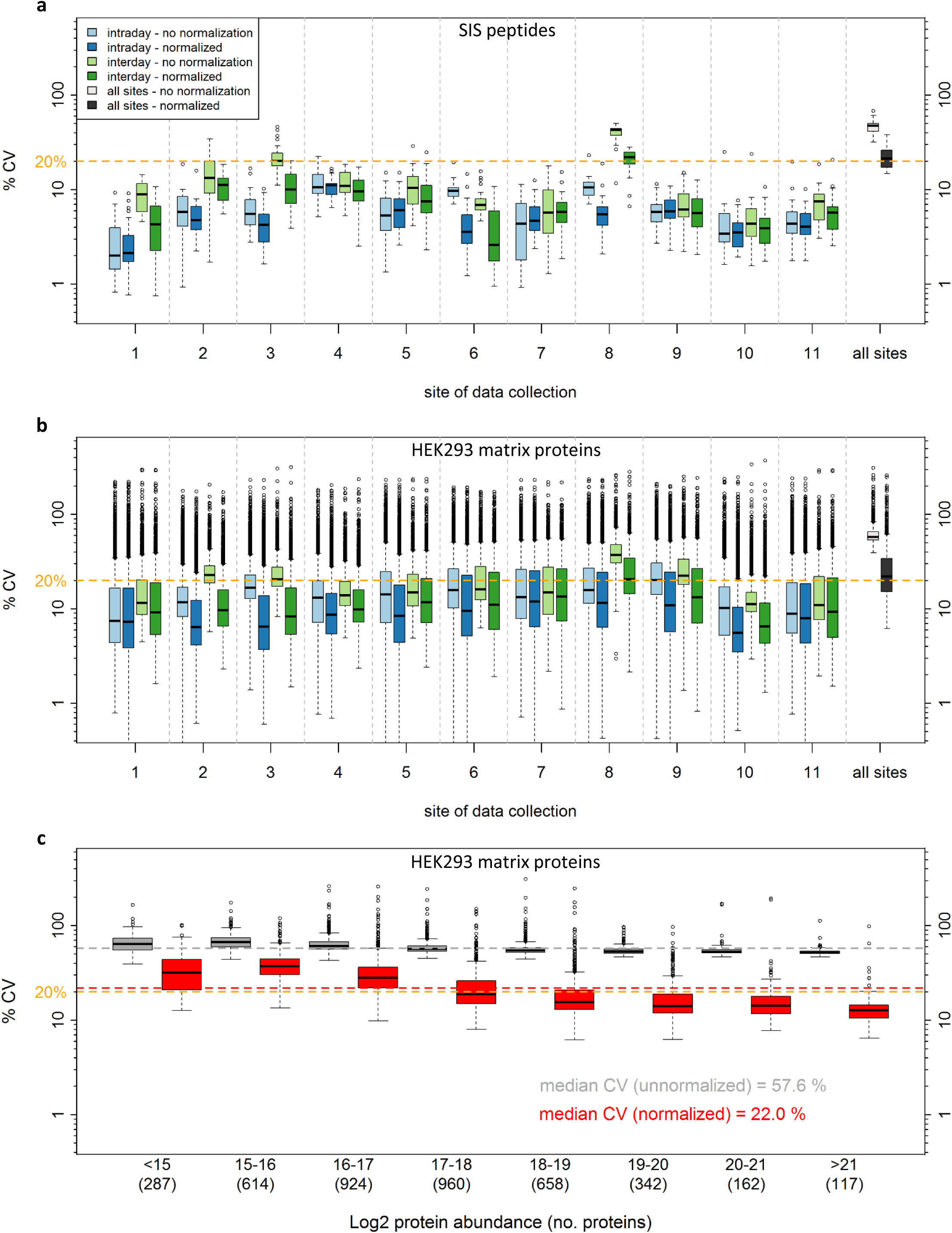
Reproducibility of SWATH-MS measurements. (a) The CVs of peak areas for each of the 30 SIS peptides for S4 sample, depicted on the y-axis using logarithmic scaling, were determined at the intra-day level within the site (light blue - without normalization, dark blue with normalization), inter-day level within site (light green - without normalization, dark green with normalization), and inter-site level (i.e. over all S4 samples in the study; light grey - without normalization, dark grey - with normalization). The orange line indicates 20% CV for visual reference. (b) Similarly, the CV of protein abundances for the 4,077 proteins that were detected in > 80% all samples were computed at the intra-day level within the site, inter-day with and inter-site (i.e. all 229 samples in the study). (c) The intersite CVs were binned based on log2 protein abundance to visualize the dependence of CV on protein abundance.

We next elected to examine the CV at protein level in the HEK293 proteome across 21 SWATH-MS acquisitions at each site. Protein level abundances were inferred from the OpenSWATH results by summing the top 5 most intense fragment ion areas from the top 3 most intense peak groups per protein^42,44,56^ **(Supplementary Tables 6-7).** For proteins where <3 peak groups were detected, all the available fragments were summed. The CVs, computed from the 4,077 proteins that were detected in > 80% of all samples, at the intra-day, inter-day, and inter-site levels were 8.3 ± 16.2%, 11.9 ± 17.2%, and 22.0 ± 17.4% respectively, after peptide level median normalization **(Fig. 3b).** The inter-site protein CV as a function of protein abundance is shown in **Fig 3c.** After normalization, the inter-site median % CV for the highest abundance protein bin in Fig. 3c was 12.7 ± 10.2% and increased to 31.9 ± 15.8% for the proteins in the lowest abundance bin, with an overall median CV of 22%

### Linearity and dynamic range

To determine the linearity and dynamic range characteristics of SWATH-MS data within and across the sites we first examined the dilution series of SIS peptides in response curves generated from the MultiQuant Software extracted ion chromatogram (XIC) analysis. A representative example for a single site is shown in **Fig 4a** (remaining sites in **Supplementary Fig 13;** source data in **Supplementary Table 8).** Peak integration for the lowest concentration peptides was manually inspected to confirm correct peptide detection and that lower limits of quantitation conformed with good bioanalytical standards (<20% CV, 80-120% accuracy, and S/N>20 at the lower limit of quantitation LLOQ^57^).

To obtain an overview of the linearity and dynamic range between sites, we computed the average peak area after summing fragment ion peak areas (unnormalized) of the SIS peptides at a given concentration point and plotted this as averaged response curve for each site **(Fig 4b and Supplementary Table 9).** The linear regressions for the average peptide area curve for each site was computed and the R^2^ values averaged 0.97. There was signal saturation for the highest concentration point (10,000 fmol), and removal of that point increased the R^2^ to 0.99. Since this study was performed, a newer instrument platform (TripleTOF 6600) has increased linear dynamic range through a different detection system, and signal saturation at high peptide load would be significantly reduced in this case. The average response curves were very similar between sites, all exceeding 4.45 orders of linear dynamic range including all data points, with an average across sites of 4.6 **(Supplementary Table 9).**

**Figure 4:**
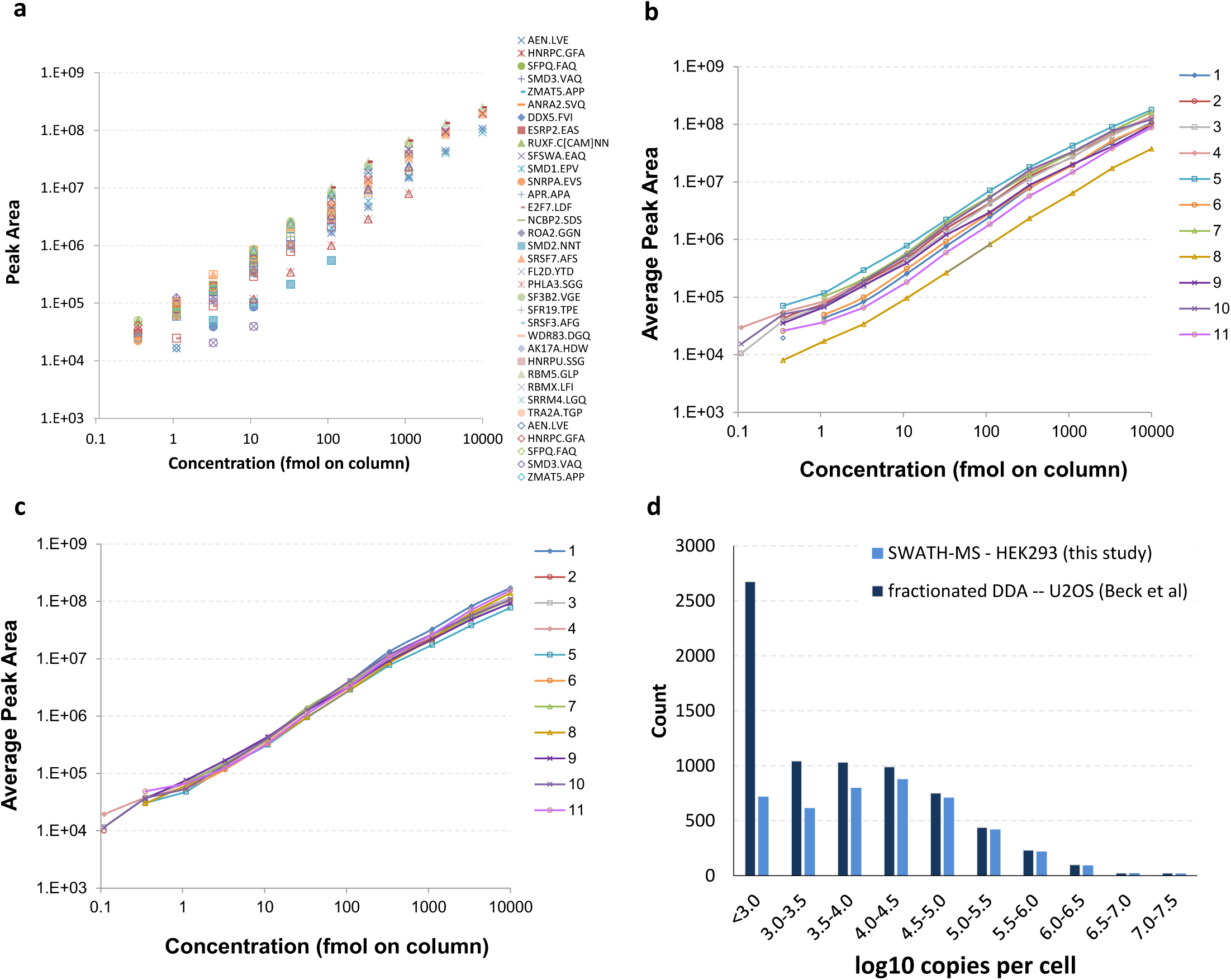
Dynamic range and linearity. (a) The response curves for each of the 30 SIS peptides for Site 1 were determined and plotted together (corresponding plots for all other sites are shown in **Supplementary Figure 13).** (b) From this data, an average response curve for each site was constructed by averaging the responses of peptides at the same concentration point. This visualization facilitates comparison of both the dynamic range and average response between sites. (c) The average response curves from (b) replotted after the normalization has been applied. (d) The proteins detected in the SWATH-MS analysis of the HEK293 proteome matrix were mapped onto a previous in-depth DDA analysis of the U20S cell line that employed multi-level fractionation to achieve deep proteome coverage. To demonstrate the dynamic range achieved by the single-shot SWATH-MS analysis we plotted the proteins detected by SWATH-MS binned by the protein copies per cell value (logl0 scale) determined from the in-depth U20S DDA study^59^. In the range 10^5-^10^7^ copies per cell the proteome coverage is essentially complete and decreases with lower copies per cell bins.

By applying the average peak area and average response curves, the data showed that the linearity and dynamic range for each site is qualitatively similar in terms of slope and span. The raw peak areas obtained from each site, however, are offset by a fixed amount across the dynamic range. When the same averaged response curve plot was constructed from values normalized based on the HEK293 proteome background, the response curves were well overlaid **(Fig. 4c).** The peptide peak area fold change between dilution steps averaged 2.66 across the concentration range, reflecting the 3-fold dilution series (ratios in the middle of the linear dynamic range are close to 3 with some compression^58^ of the ratio at the lowest and highest concentration points - **Supplementary Fig. 14).** The mean fold change for expected ratios of 9-fold and 27-fold were 7.49 and 19.6 respectively. The ratio compression is partly explained by the high peptide loads (low pmol on column range) used at the upper end of the dilution series, higher than are commonly used for this experiment type, which caused some MS signal saturation.

We next attempted to assess the dynamic range of the measurements at the protein level in the HEK293 proteome. At protein the level, no internal standard was available on which to judge dynamic range. Therefore, as a surrogate measure, we mapped the set of proteins detected in our experiment onto a previous in-depth proteomic characterization of U20S cells which estimated the copy numbers of proteins per cell^59^. Although the reference data is from a different cell line, an in-depth quantitative comparison of these two cell lines has shown that the protein abundances are well correlated (Pearson correlation ~0.8)^60^ making this a reasonable surrogate measure. From this data we can estimate that the set of proteins detected by SWATH-MS in the HEK293 cell proteome spans ~4.5 orders of magnitude, with the upper ~2.5 orders of magnitude being highly complete **(Fig. 4d).**

### Sensitivity in SWATH-MS and MS1

To get a broad view of the lower limit of quantitation (LLOQ) across the study, we plotted the percentage of the 30 SIS peptides which were reliably detected at each concentration in the dilution series from each site **(Fig. 5a; Supplementary Table 10).** Interestingly, the curves depicting % detection of peptides for the SWATH-MS data across different sites of data collection are uniform, indicating that consistent sensitivity can be achieved at different sites despite the high complexity background. The LLOQ for SWATH-MS data spanned the mid-attomole to low-femtomole range. Despite the higher complexity background proteome used in this study, the results are in good agreement with data previously obtained^32,61^.

**Figure 5:**
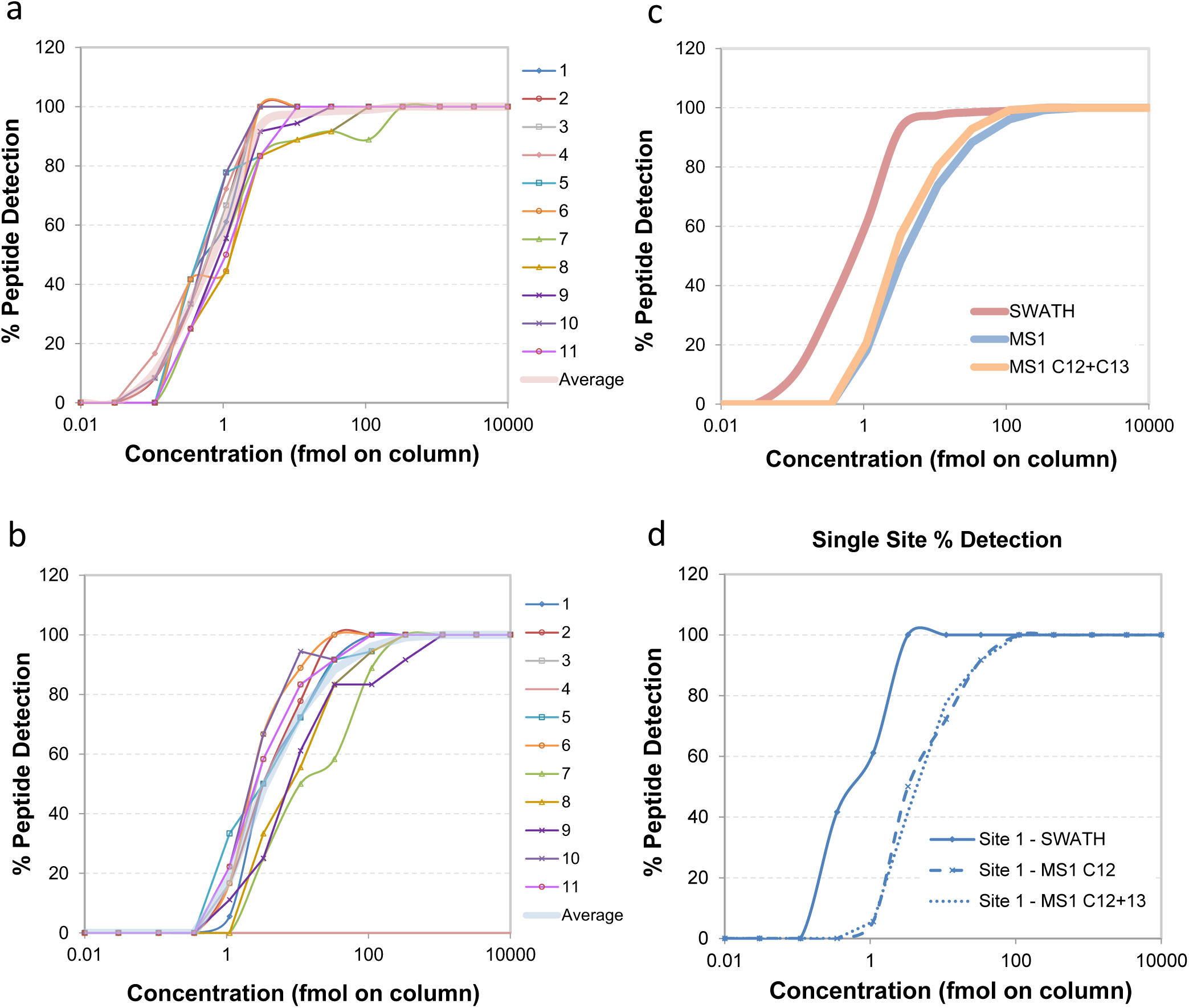
Lower limit of quantification in SWATH-MS and MSI. The percentage of the 30 SIS peptides detected at each concentration in the dilution series from each site of data collection was plotted at the SWATH-MS level (a) and the MS1 level (b). Lower limit of quantification was defined as <20% CV, S/N>20, 80-120% accuracy using linear fit with 1/x weighting in the response curve. Spectral peak widths for XIC generation were 0.02 *m/z* for MS1 and 0.05 *m/z* for SWATH-MS2, and the nominal resolving power was 30,000 and 15,000 respectively. (c) The average % detection at each concentration for all sites was determined (bold line in (a) and (b)) and overlaid to summarize detection differences between SWATH-MS and MS1 data. For the MS1 data, the C12 and C13 XIC data was also summed for comparison. (d) The data from a single site (site 1) is also shown for comparison.

As the SWATH-MS acquisition method also contains an MS1 scan in every cycle, we were able to extract XICs at the MS1 level and determine the LLOQ in MS1 mode using similar criteria as for the SWATH-MS data **(Fig. 5b).** Average lines were computed for each mode of quantification and plotted together for easy visualization **(Fig. 5c).** In our dataset the LLOQ of peptides using SWATH-MS2 quantification is nearly 1 order of magnitude lower than in MS1. The benefit in this case is explained in terms of selectivity but not absolute signal abundances. While the signal intensity of the precursor in MS1 is typically higher than the fragment ions from the SWATH-MS signal, the MS1 XICs become contaminated with interfering signals as the LLOQ is approached, whereas the SWATH-MS signal generally has less interference at lower analyte concentrations **(Supplementary Fig. 15-17; Supplementary Note 3).** This difference between SWATH-MS and MS1 level sensitivity has also been previously reported^32,35^, although usually with smaller differences between MS1 and SWATH-MS LLOQs that may be explained by the higher complexity of the sample matrix in this study. Additionally, when compared to the SWATH-MS result, the MS1 data yielded a more divergent detection rate at each concentration across sites, demonstrating that MS1 profiling has a less consistent sensitivity between labs. SWATH-MS demonstrated improved intra-lab reproducibility compared with MS1 with CV values of 8.8% and 13.2% respectively **(Supplementary Fig. 18).**

### Global similarity of quantitative protein abundance profiles

Finally, we elected to examine the global similarity of the normalized quantitative protein abundances determined by SWATH-MS across the different sites of data collection. We performed a hierarchical clustering of the study-wide log2 protein abundance matrix and plotted the resulting dendrogram in Fig. 6a. The data broadly clusters by site of data collection, whereas the day of data collection within one site generally does not cluster. To determine the similarity of the protein abundance profiles more quantitatively, we computed a pairwise Pearson correlation matrix based on the normalized log2 protein abundances of the common proteins from each pair of runs **(Fig. 6b).** The median Pearson correlation of log2 protein abundances across the entire dataset was 0.940. On average, the median Pearson correlation within a given site of data collection was only slightly higher at 0.971 (the range of site medians was 0.948-0.984). The minimum pairwise Pearson correlation between any two of the 229 files across the study was 0.868. From the above analyses, we can conclude that the quantitative similarity within sites of data collection is only marginally higher than between sites of data collection.

**Figure 6:**
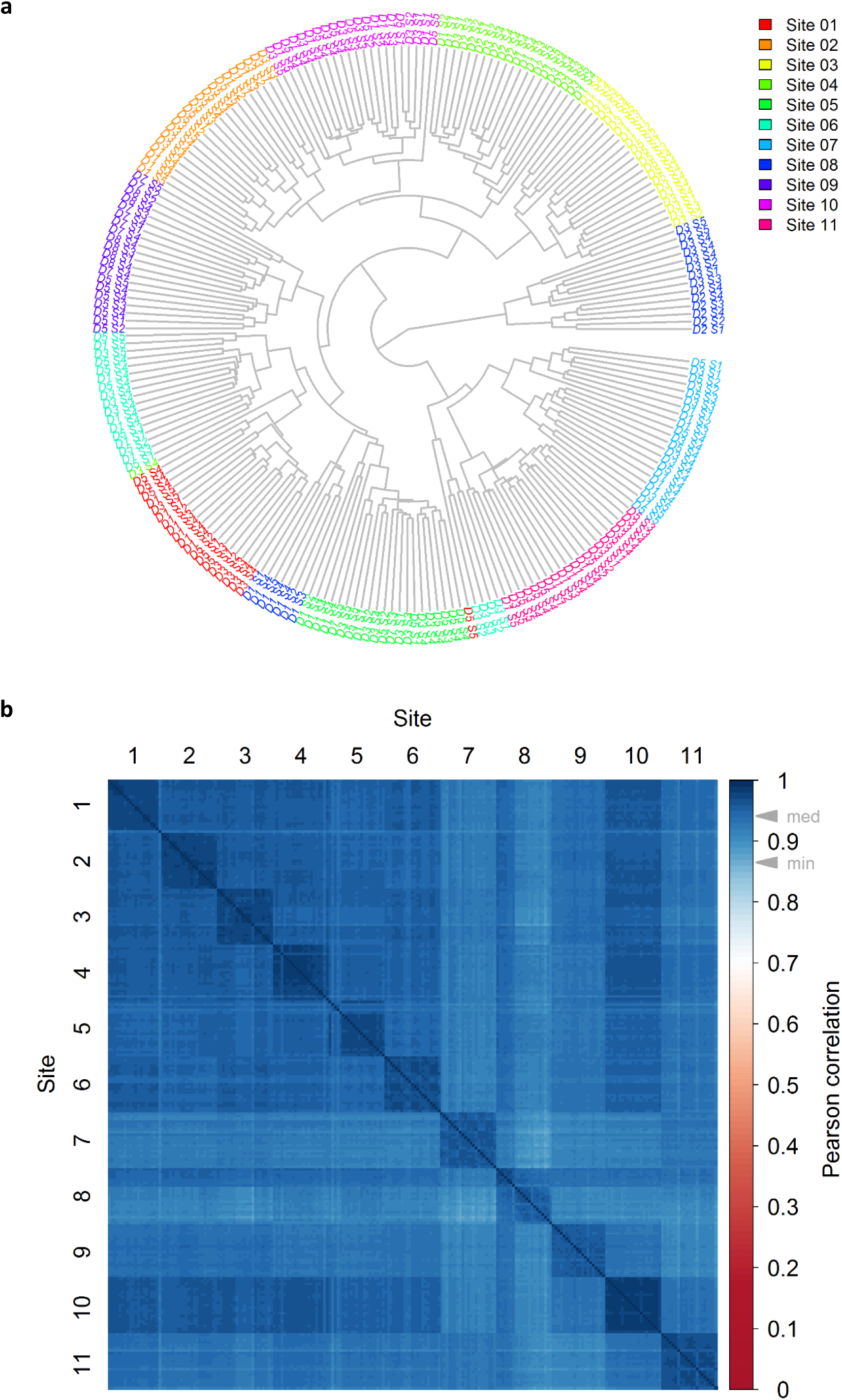
Clustering and correlation of SWATH-MS quantitative proteomes. (a) The dendrogram for the 229 samples from all sites resulting from hierarchical clustering based on the log2 protein abundances generated from the SWATH-MS data is shown. The sites are color coded as per the legend. The ‘D’ and ‘S’ notation refers to the day and sample number respectively (see Fig 1a). The samples primarily cluster by site of data acquisition whereas the day of data acquisition with one site is generally not clustered. (b) A correlation matrix showing Pearson coefficients between the 229 samples (all versus all) is shown. The samples are ordered first by site and then chronologically. The color-scale indicates the magnitude of the Pearson correlation coefficient and the grey arrowheads on the color-scale indicate the median and minimum Pearson correlation across all binary comparisons.

## Discussion

The importance of quantitative proteomics in clinical and basic research is expanding rapidly because proteins provide a direct insight into the biochemical state of the cell. In response to this need a range of MS proteomic methods have been developed. To determine the utility of particular proteomic technologies a thorough and objective assessment of their performance profiles is essential. For the widespread application of the technology, robustness, reproducibility, quantitative accuracy, data comprehensiveness and completeness are critically important performance parameters^62^. Targeted proteomics via SRM is a proven technology receiving high grades with respect to these metrics. The Clinical Proteomic Technologies for Cancer Initiative as part of the Clinical Proteomic Tumor Analysis Consortium (CPTAC) projects^26,28,29^ have demonstrated that the robust application of SRM across different labs is achievable and an Atlas of SRM assays for the entire human proteome has been published^22^. These results suggest that distributed studies with hundreds to thousands of samples and data integration between labs have become feasible. They also generally increased the confidence that smaller and larger scale, comparative proteomic studies are a reality. However, the feasibility of larger scale sample comparisons on protein numbers which exceed that quantifiable by SRM by orders of magnitude has not been demonstrated. SWATH-MS is a technique that has the potential to achieve this ambitious objective. The goal of our study was to characterize the performance of SWATH-MS acquisition across different laboratories.

The dataset analyzed in this study supports a number of conclusions relating to the above stated questions. Firstly, the set of proteins we detected across all sites is very similar and is effectively saturated after a small number of files are analyzed. This indicates that the level of data completeness from a protein quantification perspective is very high, a quality which is desirable in comparative studies. Notably, the spectral library and peptide query parameters we used to perform the analysis of the SWATH-MS data were previously published^48^ and built by a single lab independent of the current study, illustrating the generic applicability of such spectral libraries. Appropriate FDR control was key to achieving this result. Extending the FDR control to the global context (computed over all files in the analysis), in addition to extending the FDR control from the peptide query to the protein level, were critical in the project where large numbers of samples were analyzed using a large number of peptide queries. These issues are quite well understood from the perspective of DDA data analysis^55,63^ but until now have not been directly addressed for peptide-centric analysis of DIA data at large-scale. FDR control remains a challenging issue, even in DDA analysis. For example, the results from recently published drafts of the human proteome^64,65^ have been widely criticized^66,67^ for not applying protein-level FDR control, and reanalysis of one of these datasets by the original authors significantly reduced the number of reported proteins when protein-level FDR was applied^68^. An accompanying perspective discusses these issues relating to FDR control in DIA data in detail^54^.

We expect that a DDA based study could not achieve such a high level of completeness across labs due to stochastic MS2 sampling^7^ and such a study is likely to experience difficulty aligning MS1 signals arising from different labs where chromatography will inevitably vary **(Supplementary Fig. 4).** Importantly, our analysis method did not employ any alignment or propagation of peptide identifications as is commonly used in MS1 quantification from DDA data, however, we anticipate that data completeness might be further improved using a feature alignment strategy recently developed for SWATH-MS^69^. Secondly, the quantitative characteristics in terms of reproducibility, limit of detection, and linear dynamic range were also highly comparable across the data from all sites. Again, with regard to large-scale proteome quantification (i.e. 4000+ proteins) across laboratories in >200 measurements, these findings are unprecedented and have evolved to a level where many of the previously described limitations of MS-based proteomics^62^ are being significantly overcome.

In the course of analyzing the data, some interesting characteristics of SWATH-MS data became apparent. For example, one observation relates to the absolute signal response of instruments from various sites which, as expected, was variable. Interestingly, the slope, linearity and dynamic range of the response curves from the SIS peptide dilution series are essentially uniform across sites with only an offset in the intensity (y-axis) dimension differing **(Fig. 4c).** Further, the number of proteins detected at a given site was only moderately correlated with signal intensity **(Supplementary Fig. 8).** This suggests that the absolute signal intensity is not the critical metric in determining the data quality, but probably rather the signal-to-noise ratio. These observations have important consequences for normalization of label-free quantitative data and, in our study, facilitated the use of a simple global median normalization based on all of the available peptide signals from the HEK293 background proteome to effectively make the data comparable without the use of internal standards. Here, we highlight an advantage of SWATH-MS data i.e. as with MS1/DDA based quantification and, unlike more classical targeted methods such as SRM or PRM, there are very large numbers of peptides available for global normalization procedures. This dataset may also be useful for future optimization of certain general data analysis parameters, such as, selection of the most appropriate peptides for protein quantification. In this study, we used a simple method to infer protein abundance^44^, however, more advanced methods that take into account which peptides are most robust for quantification (‘quantotypic’^70^) across the study could be developed based on our data.

Another comparison that was directly possible in our dataset was that of LLOQ in either SWATH-MS or MS1 mode using XIC based analysis within the same data files. As previously reported, we found a clear benefit in sensitivity when extracting quantitative information from SWATH-MS data over MS1 data. This difference was maintained across all sites where the data was acquired, and seems to be generalizable at least with respect to the instrument setup used in this study. It should be stressed that this effect may be somewhat platform dependent, as mass analyzers with higher resolving power for MS1 spectra would facilitate smaller XIC widths, reducing interferences to some degree. This finding also allows us to speculate about the future developments in this area. As instrument scan speed continues to increase, the limitations associated with stochastic sampling of DDA may diminish^71^. However, based on the results of our comparison between MS1 and SWATH-MS quantification, we suggest that there would still be significant advantages to performing peptide quantification using MS2 signals acquired in DIA mode. Further, SWATH-MS does not necessarily rely on detection of precursor signals which can be below the detection limit where MS2 signals are still detectable and quantifiable^32,72^.

Finally, a further comparison with CPTAC and associated projects focused on targeted proteomics via SRM is of interest as it represents the most advanced work on the robustness and transferability of quantitative proteomics methods to date^26,28,29^ CPTAC has also published inter-lab studies focused on DDA analysis. However, these have primarily focused on the repeatability of peptide/protein identifications or the establishment of quality control metrics^7,10^, or on higher level similarity of differential expression analysis when different instruments and quantitative approaches were applied^66^, but have not addressed specific comparisons of quantitative metrics such as CV, LLOQ, linearity, or dynamic range. Our study is conceptually related to what was achieved by the CPTAC SRM studies although there are also some major differences. Firstly, the scope of the CPTAC SRM studies was different and included variables such as sample preparation, system suitability, and instrumentation from different vendors. In the case of our study, the decision to include only a single instrument type and model was primarily to limit the number of experimental parameters varied and, secondly, because at the outset of the project (Sept. 2013) the adoption of SWATH-type DIA analysis on other platforms was limited. As such, in our study, the main variable tested was the site of data acquisition to assess, for the first time, inter-laboratory SWATH data quality and reproducibility. In addition, CPTAC SRM studies were focused on achieving essentially clinical-grade assays^21^ for relatively discrete sets of targets, whereas our focus was on quantifying large numbers of proteins in a workflow which might be used either in a discovery mode for hypothesis generation, or in a verification mode to test large numbers of protein analytes in large cohorts. Lastly, as CPTAC has been focused on a relatively discrete set of targets it was possible to include isotope-labelled standards, which helped to determine absolute concentrations and to control matrix interference effects, whereas our study focused on label-free analysis. With these differences stated, we can suggest that our studies lead to a conceptually similar conclusion, albeit with different scopes. That is, using either targeted MS (i.e. SRM) to study discrete panels of proteins with highly validated assays, or using DIA (i.e. SWATH-MS) to study large numbers of proteins in exploratory/verification analyses, we can quantify proteins in a robust and complete manner.

This study has demonstrated for the first time that large-scale quantification of several thousand proteins is feasible with reproducible and comparable results across multiple labs. The consistent results obtained were facilitated by using the SWATH-MS approach that combines DIA with targeted data extraction, a peptide-centric scoring method, to achieve a high level of data completeness with favorable quantitative characteristics. The outcomes of this study serve to open the door for large-scale distributed quantitative proteomics studies involving hundreds to thousands of samples. The result of our study is paralleled by concurrent improvements in the robustness of data analysis tools^39^, methods for error rate control^54^, and sample preparation techniques^43^ which, collectively, advance the reproducibility and transparency of SWATH-MS. As comparative quantitative analysis of a large number of proteomes becomes accessible, we can expect to see research applications where the analysis of large numbers of samples is a prerequisite. For example, the distributed analyses of clinical material from large patient cohorts (e.g. biomarkers, precision and personalized medicine), association of protein abundances to genomic features using genetic reference collections or wild-type populations (e.g. quantitative trail locus or genome wide association studies), or large-scale perturbation screens using *in vitro* model systems (e.g. drug screens or cue-signal-response analyses) are now feasible. More broadly, the data presented here demonstrate a significant advance in the robustness of large-scale quantitative proteomics, and we expect the results from this study to increase confidence in SWATH-MS as a reproducible quantification method in life science research.

## Acknowledgements

We thank Alex Ebhardt for providing the SIS peptides for this study; Eric Deutsch for facilitating FTP data exchange; Isabell Bludau for discussions on FDR control; Uwe Schmitt for development of the PyProphet extension; Hannes Röst for discussions on normalization and data analysis; Emanual Schmid for assistance with data management. We thank Asa Wahlander and Bernd Roschitzki from the Functional Genomics Center Zurich (FGCZ) for instrument maintenance and support with the MS measurements. A-CG is the Canada Research Chair in Functional Proteomics and the Lea Reichmann Chair in Cancer Proteomics. We acknowledge funding from the Government of Canada through Genome Canada and Ontario Genomics (OGI-088, OGI-097) and Canadian Institutes of Health Research (FDN-143301) to A-CG; the National Cancer Institute Clinical Proteomics Tumor Analysis Consortium (CPTAC) grant U24CA160036 to DWC and HZ; Chinese National Basic Research Programs (2014CBA02002, 2014CBA02005). NS is supported by funding from the European Union's Seventh Framework Program HEALTH-F4-2013-602156. We acknowledge support from the NIH shared instrumentation grant for the TripleTOF system at the Buck Institute (1S10 OD016281, B.W.G.). MPM acknowledges support from the Australian Government's National Collaborative Research Infrastructure Scheme. This work was funded in part by National Institutes of Health Grant RC2 HG005805 from the National Human Genome Research Institute (NHGRI) through the American Recovery and Reinvestment Act and Grants from the National Institute of General Medical Sciences (NIGMS) grants R01GM087221, S10RR027584 and 2P50GM076547 to the Center for Systems Biology, the National Science Foundation grant MCB-1330912, AMED-CREST from Japan Agency for Medical Research and Development, and the Funding Program for Next Generation World-Leading Researchers by the Cabinet Office to MHK and SO. BCC was supported by a Swiss National Science Foundation Ambizione grant (PZ00P3_161435). RA was supported by ERC Proteomics v3.0 (AdG-233226 Proteomics v.3.0) and AdG-670821 Proteomics 4D), the PhosphonetX project of SystemsX.ch and the Swiss National Science Foundation (SNSF) grant number: 31003A_166435

## Author contributions

BCC, CLH, and YL prepared the samples, analysed the data, and wrote the manuscript. BS contributed to protocol preparation and manuscript writing. GR assisted with data analysis. BCC, CLH, YL, BS, SLB, DWC, BWG, A-CG, JMH, MH-K, GH, CK, BL, LL, SL, MPM, RLM, SO, RS, NS, SNT, SCT, and HZ acquired the data and contributed to manuscript writing. BCC, CLH, YL and RA designed and directed the study.

## Online methods

### Generation and distribution of a benchmarking sample

HEK293 cells (ATCC) were cultured in DMEM (10% FCS, 50 μg/mL penicillin, 50 μg/mL streptomycin). Cell pellets were lysed on ice by using a lysis buffer containing 8 M urea (EuroBio), 40 mM Tris-base (Sigma-Aldrich), 10 mM DTT (AppliChem) and complete protease inhibitor cocktail (Roche). The mixture was sonicated at 4 °C for 5 mins using a VialTweeter device (Hielscher-Ultrasound Technology) at the highest setting and centrifuged at 21130 g, 4 °C for 1 hr to remove the insoluble material. The supernatant protein mixtures were transferred and the protein amount was determined with a Bradford assay (Bio-Rad). Then 5 volumes of precooled precipitation solution containing 50% acetone, 50% ethanol, and 0.1% acetic acid were added to the protein mixture and kept at −20 °C overnight. The mixture was centrifuged at 20,400 g for 40 min. The pellets were further washed with 100% acetone and 70% ethanol with centrifugation at 20,400 g for 40 min. Aliquots of 2 mg protein mixtures were reduced by 5 tris(carboxyethyl)phosphine (Sigma-Aldrich) and alkylated by 30 mM iodoacetamide (Sigma-Aldrich). The samples were then digested with sequencing-grade porcine trypsin (Promega) at a protease/protein ratio of 1:50 overnight at 37 °C in 100 mM NH4HCO3^75^. Digests were combined together and purified with Sep-Pak C18 Vac Cartridge (Waters). The peptide amount was determined by using Nanodrop ND-1000 (Thermo Scientific). An aliquot of retention time calibration peptides from an iRT-Kit (Biognosys) was spiked into the sample at a ratio of 1:20 or 1:25 (v/v) to correct relative retention times between acquisitions^76^.

Thirty heavy labeled synthetic peptides^47^ were selected and the MS response for each peptide was measured. The peptides were ranked by MS response and assigned to 5 groups (A-E) to ensure there was a range of responses across in each group. These peptides groups were diluted into the matrix described above across a concentration range to create the 5 different samples to be analyzed **(Fig 1a; Supplementary Table 1 and 2).** Finally, samples were shipped on dry ice to the 11 sites.

### SWATH-MS measurements

Peptide mixtures were separated using reversed phase nanoLC using either a nanoLC Ultra system or a nanoLC™ 425 system (SCIEX). Most sites (9 of 11) used a cHiPLC^®^ system (SCIEX) operated in serial column mode (for detailed acquisition information please see SOP in **Supplementary Protocol 1),** fitted with two cHiPLC^®^ columns (75 <m × 15 cm ChromXP™ C18-CL, 3 μm, 300 Å) to give a total column bed length of 30 cm (Site configuration details in **Supplementary Table 11).** Two sites used PicoFrit emitter (New Objective) packed to 30 cm with Magic C18 AQ 3 μm 200 Å stationary phase. Peptide samples (2 <L injection) were first loaded on the first cHiPLC column and washed for 30 mins at 0.5 μL/min using mobile phase A (2% acetonitrile in 0.1% formic acid). Then, elution gradients of approximately 5-30% of mobile phase B (98% acetonitrile in 0.1% formic acid) in 120 mins were used to elute peptides off the first column and through the second cHiPLC column. Both columns were maintained at 35 °C for retention time stability. Similar separations were performed across all sites. Gradients were allowed to minimally vary from site to site to obtain similar peptide separations (see **Supplementary Table 11** for gradient information).

Eluent from the column was introduced to the MS system using the NanoSpray^®^ Source into a TripleTOF 5600 system with Analyst^®^ Software TF 1.6 (SCIEX) and the variable window acquisition beta patch. The SWATH-MS acquisition methods were built using the SWATH-MS Acquisition method editor and a pre-defined variable window width strategy using 64 windows **(Supplementary Table 13).** The Q1 mass range interrogated was 400-1200 *m/z,* and MS2 spectra were collected from 100 - 1500 *m/z* with an accumulation time of 45 msec per variable width SWATH window. A TOF MS scan (250 msec, 400 - 1250 *m/z)* was acquired in every cycle for a total cycle time of ~3.2 sec. Nominal resolving power for MS1 and SWATH-MS2 scans were 30,000 and 15,000 respectively. The collision energy curve was controlled across all instruments (CE = 0.0625 * *m/z* −3) and the collision energy spread was defined in the variable window table **(Supplementary Table 13).** The acquisition order is outlined in the **Supplementary Table 14.**

### Pilot phase quality control assessment

SWATH-MS acquisition data from the pilot study phase were processed using the SWATH^®^ Acquisition MicroApp 2.0 in PeakView^®^ Software 2.2. A previously published proteome library containing mass spectrometric coordinates for 10,000+ human proteins^48^ was used for data processing. iRT standard peptides (Biognosys) were included in the library for automatic retention time calibration of each different sample set with the ion library retention times. Peak group detections were filtered at a 1% global FDR and metrics were compared using Excel (this corresponds to data in **Supplementary Fig. 1** only).

### Automated analysis of SWATH-MS data for human HEK293 proteome matrix

The SWATH-MS data analysis was performed using OpenSWATH (OpenMS v2.0) essentially as described^35^ except that the improved single executable OpenSwathWorkflow was used instead of the multi-step workflow to perform peak-picking and feature detection and the following parameters were changed: *m/z* extraction window = 75 ppm, RT extraction window = 900 seconds. The spectral library used as input for peptide queries in the OpenSWATH analysis was a previously published proteome library containing mass spectrometric coordinates for 10,000+ human proteins built by combining several hundred DDA analyses of various human cell and tissues types^48^.

Semi-supervised learning to optimally combine OpenSWATH peptide query scores into a single discriminant score, and q-value^49^ estimation to facilitate FDR control, were performed using an extended version of PyProphet^77^ (PyProphet-cli v0.19 - https://github.com/PyProphet). PyProphet was run both using the experiment-wide context (local-global option in PyProphet - q-values are generated for every peptide query and protein in every sample) and the global context (global-global option - only one q-value for every peptide query and protein representing the highest scoring instance over the whole experiment), with a fixed λ of 0.4. The set of peptide peak groups used for learning the score weights of OpenSWATH sub-scores to produce a single discriminant score were sampled with a ratio ≈ 1/(no. of samples) in the analysis (for aggregated analysis of all sites 0.005, and for analysis of individual sites 0.05). The sets of peak groups detected at 1% FDR and proteins detected at 1% FDR in the global context were used as a filter to restrict the set of peak groups and proteins in the experiment-wide context. The filtered table from the experiment-wide context was then filtered at 1% FDR at the peptide query level. A protein was considered as detected in a given sample if it passed these consecutive filters (see **Supplementary Note 2** for further discussion on FDR control).

Normalization was achieved by equalizing medians at the peak group level. The normalization coefficients derived from the peak groups in HEK293 matrix were also used to normalize the peak areas determined by MultiQuant analysis (below) of the SIS peptides. Protein abundances were inferred by summing the top 5 most intense fragment ion peak areas from the top 3 most intense peak groups using the aLFQ software^56^ (v1.33). Where < 3 peak groups were detected, the available peak groups were summed. Coefficients of variation (% CV) were computed as 100*standard deviation/mean. Hierarchical clustering was performed using the dist and hclust functions in R (v3.2.2) using log2 transformed protein abundances and visualized using the R package ape (v3.3). Pearson correlation coefficients were computed using the R package Hmisc (v3.17) and visualized using the R package corrplot (v0.73).

### Analysis of SWATH-MS data for 30 SIS peptides in dilutions series

The SWATH Acquisition data obtained from all sites was processed using MultiQuant Software 3.0. The same quantification method **(Supplementary Table 15)** was used across all sites and consisted of 3-4 fragment ion extracted ion chromatograms (XICs) extracted and summed together to produce a peptide area. Spectral peak widths for XIC generation were 0.05 for MS2 and 0.02 for MS. Peak integration was done using the MQ4 algorithm. The curve for each peptide was evaluated and the LLOQ was determined in accordance with bioanalytical standards^49^ (<20% CV, S/N>20, 80-120% accuracy; linear fit with 1/x weighting). A number of analytical aspects were evaluated, including the reproducibility of the peptide peak areas, the lower limit of quantitation (LLOQ) for each peptide, the signal/noise ratios using the relative noise approach in the MultiQuant Software, and the reproducibility and accuracy of the concentration.

